# Degradation of biopesticidal triterpenoid saponins by the soil bacterium *Arthrobacter* sp. α-11c

**DOI:** 10.1101/2025.08.18.670823

**Authors:** Kitzia Yashvelt Molina-Zamudio, Malbor Dervishi, Chen Wang, Marco Rami, Patrick Denis Browne, Hans Christian Bruun Hansen, Søren Bak, Mette Haubjerg Nicolaisen

**Author notes:** **Corresponding author:** Mette Haubjerg Nicolaisen. Authors contributed equally (Kitzia Yashvelt Molina-Zamudio, Malbor Dervishi). **Co-authors:**.

## Abstract

**Background:** Saponins, a diverse group of glycosylated triterpenoid and steroid compounds produced by plants, exhibit potent insecticidal activity and are promising candidates for sustainable pest management. However, their potential persistence in soil raises concerns about ecological impacts, highlighting the need to understand microbial degradation rates and pathways.

**Results:** This study reports the isolation of *Arthrobacter* sp. α11c, and its ability of fully degrading two hederagenin-based pentacyclic triterpenoid saponins: α-hederin, hederacoside C, and the corresponding sapogenin hederagenin. All three compounds were fully metabolized within six hours when provided as carbon source. Fractionation experiments confirmed intracellular uptake and degradation of the saponins.

Genome analysis of α11c revealed diverse and multiple glycosidase genes, yet basal glucosidase activity remained unchanged when cultured with these saponins. Comparative genomics with *Arthrobacter* sp. α12b, a closely related strain unable to degrade saponins, revealed two unique glycosidase gene clusters in α11c, suggesting a role in adaptation to saponin metabolism.

**Conclusion:** These findings enhance our understanding of bacterial degradation of plant triterpenoid saponins and provide a foundation for evaluating their environmental fate. This knowledge supports the safe and sustainable use of saponin-based biopesticides in agriculture by identifying microbial partners that contribute to their breakdown in soil ecosystems.

## Introduction

In the strive for sustainable and environmentally friendly pest management, plant derived compounds have a high but untapped potential (Zhang et al., 2023). Among these, saponins have emerged as promising candidates for biopesticide development (Jiang et al., 2019).

Saponins are amphipathic glycosides broadly occurring in over 100 plant families (Vincken et al., 2007). They are classified into three main categories based on the sapogenin structure; triterpenoid saponins, which are typically pentacyclic, steroidal saponins that are derived from sterol metabolism and are typically tetracyclic, or steroidal glycoalkaloids that are steroidal triterpenoids conjugated with an alkaloid moiety (Cárdenas et al., 2019; Erthmann et al., 2018; Nakayasu, Ohno, et al., 2021; Nakayasu, Yamazaki, et al., 2021). Triterpenoid saponins are typically glycosyated at C3, classified as monodesmocidic, or at both C3 and C28, classified as bidesmocidic (Augustin et al., 2011).

Saponins have many biological functions and have recently gained interest as potential biopesticides against agricultural pests. Saponins may function as biopesticides through various mechanisms, including competition with sterol uptake, interference with hormone signaling, and damage to the membranes of insect guts (De Geyter et al., 2012; Hussain et al., 2019; Roopashree & Naik, 2019). Oleanane-type saponins, a subgroup of the pentacyclic triterpenoid saponins that are naturally occurring and accumulate to high levels in the leaves and fruits of *Hedera helix* (Vercruysse et al., 2023), exhibit strong cytolytic activity (Chwalek et al., 2006; Vo et al., 2017). This cytotoxic activity appears to be related to their ability to directly interact with membrane sterols, such as cholesterol and ergosterol, allowing them to form pores in eukaryotic membranes (Augustin et al., 2011; Chwalek et al., 2006; Coleman et al., 2010; Orczyk et al., 2020). The oleanane-type sapogenin hederagenin (Hed) from *Hedera helix* exhibits potent effects against the Egyptian cotton leafworm (*Spodoptera littoralis*) (Adel et al., 2000; Chen et al., 2024) and monoglucosylated oleanane-type saponins have been shown to have biopesticide activity against diamondback moth (*Plutella xylostella*) and the tobacco hornworm (*Manduca sexta*) (Liu et al., 2019). Similarly, the monodesmosidic saponin α-hederin (α-Hed), exhibit significant toxicity against *Daphnia magna* (crustacean), *Enchytraeus crypticus* (annelid worm), *Saccharomyces cerevisiae* (baker’s yeast), and *Raphidocelis subcapitata* (micro algae) (Dervishi et al., n.d.), while bidesmosidic saponins, including hederacoside C (HedC) are inactive against these organisms. Interestingly, the species that are most susceptible to α-Hed were those with animal or fungal sterols, whereas the algae, which contains phytosterols, was less susceptible (Dervishi et al., n.d.). This indicates that cytotoxicity depends on both saponin structure and the membrane sterol composition of the target (Podolak et al., 2010). This may enable the development of target-specific biopesticides, positioning saponins as promising alternatives to conventional chemical pesticides.

However, the high solubility and low sorption affinity of saponins, due to their soap-like, surface-active properties, may increase their leaching potential when used as agrochemicals, raising concerns about soil, groundwater and surface water contamination (Cao et al., 2013; Nemček & Hagarová, 2021). This raises concerns about potential environmental risks and highlights the need to assess their fate and persistence in the soil. The green transition seeks to balance effective pest control with environmental safety, and approval of (bio)pesticides requires degradation rate parameters (DegT50 values) (European Food Safety Authority, 2014). Previous studies have shown that triterpenoid saponins from *Quillaja saponaria* when applied to soil, are quickly adsorbed, especially at high clay and organic matter content (Wahab & Yusup, 2021). However, it can be difficult to assess whether degradation of saponins in soil is driven by microorganisms or just an artefact of saponins remaining attached to soil particles. Therefore, soil bacteria isolates are highly needed to study degradation rates and natural degradation processes in the soil matrix.

Studies on the degradation of saponins have mainly focused on steroidal saponins. Complete steroid degradation in bacteria has only been described in *Comamonas testosteroni* TA441, *Mycobacterium tuberculosis* H37Rv, and *Rhodococcus jostii* RHA1 (Bergstrand et al., 2016; Chiang et al., 2019). However, regardless of the sapogenin, the initial step in the degradation of saponins is expected to be by deglycosylation (Overney & Huang, 2020). *In vitro* studies have demonstrated that members of the Micrococcaceae family have glycoside hydrolases such as α-rhamnosidase, β-galactosidase, and β-glucosidase that may biotransform steroidal saponins, specifically glycoalkaloids (Hennessy et al., 2020; Wang et al., 2022) and the steroidal saponin ginsenosides (Park et al., 2017). Upon deglycosylation, the liberated steroidal sapogenin undergoes microbial degradation initiated by oxidation of the A-ring. This transformation is catalyzed by enzymes such as 3α,20β-hydroxysteroid dehydrogenase or 3-oxosteroid-1-dehydrogenase (Bergstrand et al., 2016; Chiang et al., 2019).

To date, the only reported microbial modifications of the sapogenin from oleanane-type triterpenoid saponins have been C3 oxidation and C5 hydroxylation of oleanolic acid, as well as C3 oxidation of glycyrrhetinic acid (Luchnikova et al., 2022, 2023) and betulin (Grishko et al., 2013; Maltseva et al., 2024) by cultures of *Rhodococcus rhodochrous*. However, no study has fully characterized the microbial degradation of pentacyclic triterpenoid hederagenin-based saponins and their sapogenin. Therefore, the present study aimed to isolate microbes that can degrade oleanane-type saponins, specifically the bioactive monodesmosidic triterpenoid saponin α-Hed, the non-bioactive bidesmosidic saponin HedC, along with their common hederagenin sapogenin backbone, creating a platform for studying saponin biodegradation in soil.

## Experimental Procedures

### Isolation of Strains, Culture Media, and Culture Conditions

Bacteria were isolated from a topsoil sample collected from the upper 0–5 cm layer obtained at the University of Copenhagen experimental site in Taastrup, Denmark. This soil contains 16.5% clay, 17.1% silt, 27.5% fine sand, and 27.2% coarse sand, with a near-neutral pH of 7.04 and approximately 3.32% soil organic matter by weight. To enrich bacteria capable of growing on saponins, a soil suspension was prepared by vortexing 10 g of soil in 100 mL of distilled water. The soil suspension was amended with saponins (α-Hed and HedC) (>98.0% HPLC, Chengdu Biopurify Phytochemicals Ltd.) or sapogenin (Hed) (≥97.0% HPLC, Sigma-Aldrich) at a final concentration of 20 µM and incubated for 48 hours at 20°C. After incubation, the suspension was serially diluted and plated on 1/10 Reasoner’s 2A agar (R2A) amended with saponins or sapogenin by plating 100 µL of each saponin/sapogenin (20 µM), dissolved in mineral buffer (8.5 g/L KH₂PO₄, 21.75 g/L K₂HPO₄, 33.4 g/L Na₂HPO₄·2H₂O, 0.5 g/L NH₄Cl, 27.5 g/L CaCl₂, 22.5 g/L MgSO₄·7H₂O, and 0.25 g/L FeCl₃·6H₂O, with the pH adjusted to 7.4). The plates were incubated in the dark at 20°C for 72 hours.

To obtain pure cultures, single colonies were transferred into 15 ml test tubes containing 2 ml of 1/10 R2A medium. The tubes were incubated for 72 hours at 20°C under agitation using an orbital shaker at 200 rpm in the dark. After incubation, the cultures were serially diluted and plated onto R2A medium to re-isolate single colonies. This process was repeated until pure isolates were achieved. For subsequent experiments, the isolates were prepared by streaking onto R2A agar plates and incubation for 72 hours at 20°C. A single colony from each isolate was transferred into 2 ml of 1/10 R2A broth and incubated for 72 hours at 20°C under agitation in the dark using an orbital shaker at 200 rpm. After incubation, cells were washed twice in MiliQ water and resuspended in 1 ml of sterile MilliQ water.

### Growth assay

Optical density at 600 nm (OD600nm) was used as a proxy to assess bacterial growth when cultured with α-Hed, HedC, and Hed as the main carbon source. Two mM saponin/sapogenin stock solutions were prepared in 100% ethanol to ensure solubility and stored at 4°C. Isolates precultured as described above were washed and 200 µl of washed bacterial cells were added to 1.8 ml of α-Hed, HedC, or Hed solutions, which had been previously adjusted such that the final concentration of each compound in the final mixture was 20 µM in 1% EtOH. Tubes containing 1% EtOH solution without saponins/sapogenin were used as control medium. Additionally, tubes containing α-Hed, HedC, and Hed solutions without bacteria were prepared as blank controls. The tubes were incubated at 20°C for 24 hours under agitation at 200 rpm in the dark using an orbital shaker, and bacterial growth was monitored based on OD600nm. Each of the bacterial strains were cultured in biological triplicates.

### 16S RNA gene sequencing and phylogenetic analysis

Genomic DNA was extracted from overnight cultures using the alkaline lysis method (Zhang et al., 2021). PCR amplification of the 16S rRNA gene was performed in a final volume of 50 µL, comprising 2 µL of template DNA, 25 µL of NZYTaq II 2X Green Master Mix (NZYTech, Portugal), and 1 µL each of the universal primers 27F (5′-AGAGTTTGATCCTGGCTCAG-3′) and 1492R (5′-CGGTTACCTTGTTACGACTT-3′) (Galkiewicz & Kellogg, 2008), resulting in a final primer concentration of 2µM. The thermal cycling conditions involved an initial denaturation step at 97°C for 10 seconds, followed by 35 cycles of annealing at 57°C for 30 seconds, and extension at 72°C for 45 seconds. The amplicons were purified using AmPure XP magnetic beads (Beckman Coulter, US) following the manufacturer’s protocol and Sanger sequenced (Eurofins, Germany). Subsequently, a multiple sequence alignment was constructed using the ClustalW algorithm with default parameters in MEGA-X software (Kumar et al., 2018; Thompson et al., 1994). The multiple alignment was used to construct a maximum likelihood phylogenetic tree in MEGA-X, supported by 1000 bootstrap replicates (Kumar et al., 2018).

### Full genome sequencing and genome mining for glycosidases

Genome libraries of *Arthrobacter* sp. α11c, and *Pseudoarthrobacter* sp. α12b were prepared with a Nanopore rapid barcoding kit, SQK-RBK114-96, and sequenced in high accuracy mode (260 bps) in a FLO-MIN114 (R10.4.1) flow cell in a MinION Mk1B using MinKNOW version 22.10.10. Sequencing data was base called with the model’dna_r10.4.1_e8.2_260bps_sup@v4.1.0’ and demultiplexed using Dorado version 0.5.3 (https://github.com/nanoporetech/dorado). Genomes were assembled using flye (Version 2.9.3) and annotated using Prokka (Version 1.13) (Cuccuru et al., 2014; Lin et al., 2016; Seemann, 2014). The annotated genomes were search for glycosidases. Proteins annotated as glycosidases with an E.C 3.2.1 classification were filtered and extracted as a fasta file using the SeqIO module of BioPython (Cock et al., 2009). The *Arthrobacter* sp. α11c whole genome was aligned with *Pseudoarthrobacter* sp. α12b using mummer2circos (Version 1.4.2) (https://github.com/metagenlab/mummer2circos) under default conditions, where the Nucmer is the default option for alignment (Delcher et al., 2002; Krzywinski et al., 2009).

### Glucosidase activity

β-glucosidase activity was quantified using a modified version of the colorimetric method with p-nitrophenyl-β-d-glucopyranoside (pNPG) (Sigma-Aldrich, Denmark) as substrate (Chang et al., 2011; Hennessy et al., 2020; Strahsburger et al., 2017). The assays were performed in clear flat bottom 96-well plates (Thermo Scientific™) containing 150 µL of 5 mM pNPG in 200 mM sodium phosphate buffer, pH 7.0, and 50 µL of bacterial supernatant or intracellular fraction (final volume of 200 µL). The plate was incubated at 37°C for 1 hour with shaking at 200 rpm, and the reactions were stopped by addition of 200 µL of 2 M Na_2_CO_3_. Absorbance was measured using a FLUOstar Omega Microplate Reader (BMG LABTECH) at 405 nm.

### Saponin intra-and extracellular biodegradation assay

To assess α-Hed, HedC, and Hed degradation by *Arthrobacter* sp. α11c, we performed an experiment under the same culture conditions as the used for the growth assay. After 24 h incubation, the samples were collected to analyze the amount of saponins/sapogenin accumulating intracellularly and extracellularly.

To collect the extracellular culture fractions, the *Arthrobacter* α-11c culture was centrifuged at 30000 x g for 5 minutes, and then 500 µl of the supernatant was collected for LC-MS and GC-MS analysis. To assess saponins/sapogenin that might have precipitated on the cells during centrifugation, the remaining supernatant was completely removed, and the cell pellet was resuspended in 1 ml of 96% ethanol. Subsequently, the cells were pelleted at 30000 x g for 5 minutes, and 500 µl of the ethanol-soluble fraction was collected for LC-MS and GC-MS analysis; this fraction was designated as cell-washed. To collect the intracellular fraction, the remaining pellet was resuspended in 1 ml of 96% ethanol and sonicated for 15 minutes using an ultrasonic water bath (Struers, Denmark). After sonication, the lysed cell suspension was centrifuged at 30000 x g for 5 minutes to remove cell debris, and then 500 µl of supernatant was collected for LC-MS and GC-MS analysis.

### Saponin analysis using LC-qToF-MS/MS

To identify the saponins/sapogenin, metabolite analysis was carried out using LC-qToF-MS/MS. The analysis was performed on a Dionex UltiMate 3000 Quaternary Rapid Separation UHPLC+ focused system (Thermo Fisher Scientific, Germering, Germany). A Kinetex 1.7 μm XB-C18 column was used for separation. The mobile phases were 0.05% (v/v) formic acid in Milli-Q water (Phase A) and acetonitrile with 0.05% (v/v) formic acid (Phase B). The gradient program was as follows: 0.0–1.0 min with 5% B; 1.0–2.0 min from 5–30% B; 2.0–14.0 min from 30–70% B; 14.0–15.0 min at 70–100% B; 15.0–16.0 min at 100% B; 16.0–17.0 min back to 5% B; and 17.0–20.0 min at 5% B. The mobile phase flow rate was set at 300 μL/min, with the column temperature maintained at 30°C. The UHPLC system was coupled to a Compact micrOTOF-Q mass spectrometer (Bruker, Bremen, Germany) equipped with an electrospray ionization (ESI) source in negative ion mode. The ion spray voltage was set to-3900 V, with a dry temperature of 250°C and a dry gas flow of 8 L/min. Nitrogen was used as dry gas, nebulizer gas, and collision gas. The nebulizer gas pressure was 2.5 bar, and the collision energy was 10 eV. MS spectra were collected in the m/z range of 50 to 1400, and MS/MS spectra were recorded from 200 to 1400 m/z, with a sampling rate of 3 Hz. Samples were filtered prior to run through a 0.2 μm Durapore 96-Well plate membrane (Millipore, Ireland) and stored to 4 °C prior to measurement.

### Sapogenin analysis using GC-MS

Prior to extraction every sample was spiked with 20 μL of 5-α-cholestane (≥97.0% HPLC, Sigma-Aldrich) as an internal standard. Samples from the supernatant (extracellular fraction), cell-washed and intracellular fractions were collected and extracted three times with 1mL of hexane, pooled, and evaporated at 55°C under nitrogen flow. The samples were then redissolved in 50 μL of hexane and 30 μL were transferred to a MS vial together with 30 μL of N,O-Bis(trimethylsilyl)trifluoroacetamide (BSTFA, Sigma-Aldrich) to silylate the products. The mixture was then silylated by heating to 60°C for one hour before being injected into Gas-chromatography-mass-spectrometry (GC-MS). GCMS was carried out using a Shimadzu GCMS Nexis-2030 Ultra system (Shimadzu, Germany) equipped with an HP-5MS UI (Agilent Technologies, USA) capillary column (Length: 30.0 m; Thick: 0.25 um; Dia: 0.25 mm), using helium as carrier gas (linear velocity of 30.0 cm/s). Hexane samples were injected via spitless injection using an injection volume 2 μL and temperature 250°C. The GC program was as follows: Initial column temperature of 60°C, 0.0–1.0 min at 60°C; 1.0–8.0 min from 60–280°C; 8.0–44.0 min from 280–310°C; back to 60°C for new measurement.

### Biodegradation kinetics

To determine the degradation rate of α-Hed, HedC, and Hed, *Arthrobacter* sp. α11 cells were pre-cultured as described above, washed twice in MiliQ water, and resuspended in saponin/sapogenin solution to adjust to an OD600nm of 1.5. Two-hundred µl of the washed cells were inoculated into 1.8 ml of 20 µM solutions of α-Hed, HedC, and Hed. Inoculated test tubes were incubated at 20°C under 200 rpm agitation in the dark using an orbital shaker. Samples were collected from independent test tubes at 20, 40, 60, 180, and 360 minutes. The intra-and extracellular culture fractions were collected for LC-MS and GC-MS analysis as described for the biodegradation assay. The collected samples were kept at 4°C until GCMS/LCMS analysis.

Saponin concentrations (µM) were calculated using calibration curves based on standard curves of authentic standards (**Fig. S5**). To determine the dynamics of the saponins’ supernatant reduction, the data was fitted using the first-order degradation decay formula [Sap]=[Sap0]*exp(-k*t) from GraphPad Prism version 10.0.0 (GraphPad Software, Boston).

### Statistical analysis

The data sets collected from all the repetitions were analyzed for normality using the Shapiro-Wilk, Kolmogorov-Smirnov, and QQ plots. The data was normally distributed, and to determine the significance between treatments, we used a one-way ANOVA test, followed by Dunnett’s multiple comparison test, and the two-way ANOVA test. The analyses were conducted using GraphPad Prism version 10.0.0 (GraphPad Software, Boston).

## Results

### Isolation and taxonomic identification of isolated bacteria

A total of 30 bacterial strains were isolated from the saponin-supplemented plates. Based on full length16S rRNA gene sequencing isolates were found to belong to nine different genera i.e. *Massilia* spp., *Variovorax* spp., *Pseudomona*s spp., *Phyllobacterium* spp., *Ensifer* spp., *Sphingomonas* spp. *Paenarthrobacte*r spp., *Arthrobacter* spp., and *Pseudoarthrobacter* spp. (**Fig. S1**).

### Some bacteria can use saponins as a carbon source

To assess which bacteria could use the saponins α-Hed and HedC or the sapogenin Hed as a carbon source for growth, the 30 bacterial isolates were cultured separately on α-Hed, HedC, and Hed amended plates. Optical density (OD600nm) was used as a proxy for bacterial growth, and this identified four strains *i.e., Arthrobacter* sp. α11c, *Arthrobacter* sp. C19c, *Sphingomonas echinoides* α14c, and *Phyllobacterium* sp. C19b, which demonstrated a significant increase in OD600nm in the presence of saponins as compared to the water control during an initial screening (**Fig. S1**). An additional three *Massilia* strains had increased OD600nm values when grown on Hed. However, OD600nm was not found to be significantly higher than the control. Three of the strains showing significant increase in OD600nm values were selected together with six strains *i.e., Pseudarthrobacter* sp. Α12b, *Sphingomonas echinoides* C3c, *Pseudomonas migulae* A11a, *Variovorax* sp. A4b, and *Massillia cellulosilytica* A8b, *Massillia frigida* α11c, which did not demonstrate changes in the OD600nm, for a repetition experiment to confirm the reproducibility of the results.

The repetition of the growth assay confirmed a significant increase in OD600nm for *Phyllobacterium* sp. C19b when cultured with both saponins (*P* < 0.0001) and the sapogenin (*P* < 0.0001), for *Massilia* sp. α11d when cultured with the sapogenin (*P* < 0.01), and *Arthrobacter* sp. α-11c when cultured on α-Hed (*P* < 0.001) and HedC (*P* < 0.01) (**Fig. 1**).

**Figure 1.**
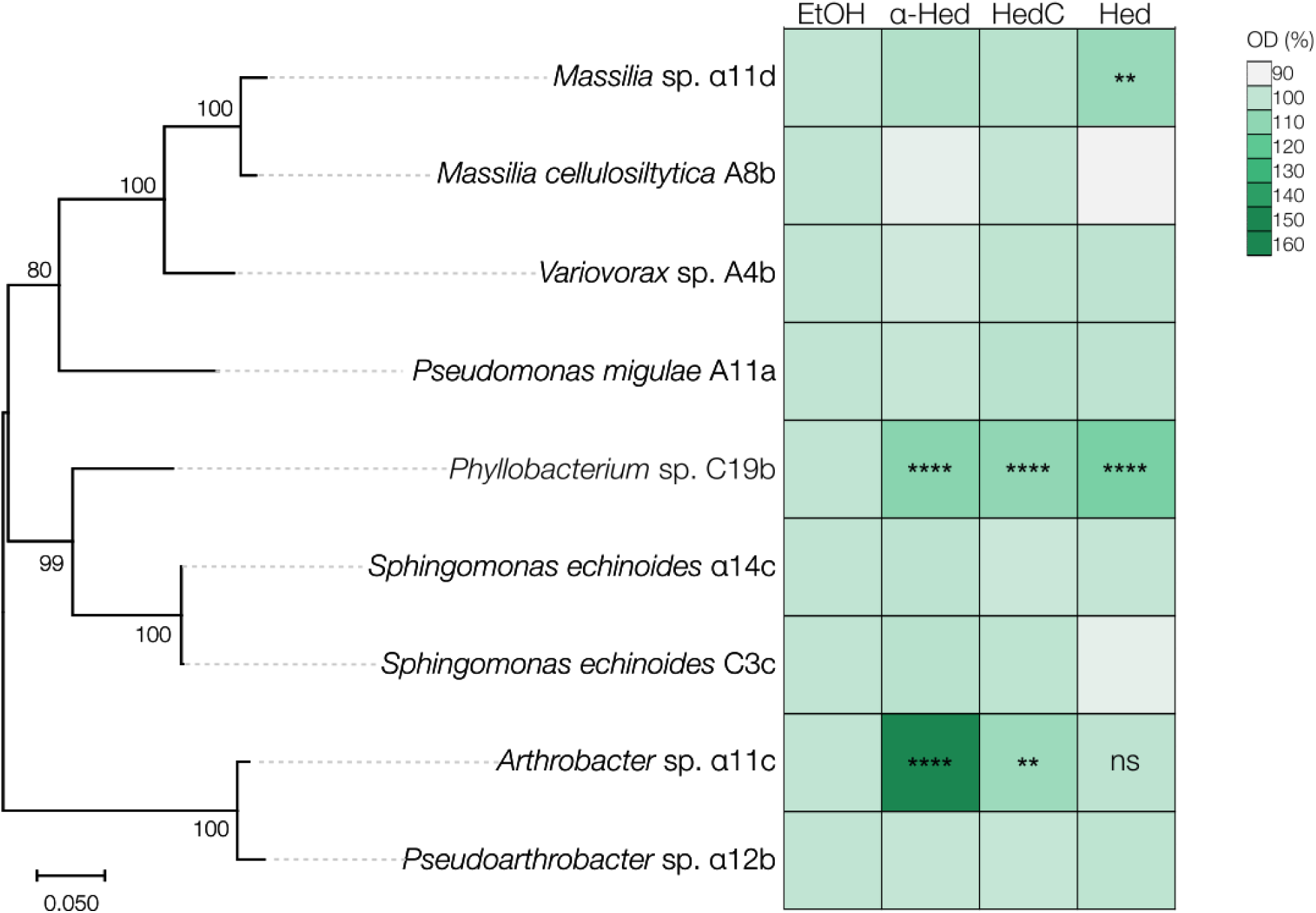
Bacteria can use saponins as a carbon source. Phylogenetic tree of the nine selected bacterial strains and corresponding OD600 values after growth in saponin-containing media, represented as percentages (%) relative to the EtOH 1% control. Based on a multiple sequence alignment of 16S rRNA gene sequences, a maximum likelihood phylogenetic tree was constructed in MEGA-X; bootstrap values (1000 iterations) are shown at the nodes. Bacterial cells were inoculated into media containing saponins (α-Hed and HedC) or the corresponding sapogenin (Hed) to a final concentration of 20 µM, each dissolved in 1% EtOH (used as control). OD600 was measured after 24 h of incubation. Statistical analysis was performed using one-way ANOVA; ns, not significant; **, P < 0.01; **, P < 0.0001. Data represents means from three biological replicates.

### Genome mining of potential saponin biodegradation genes

It is known that some bacteria can degrade and grow on saponins, accordingly we hypothesized that glycosidase activity (EC 3.2.1) is involved in degradation of the saponins α-Hed and HedC, by releasing glycosides by hydrolysis to support bacterial growth. To explore this hypothesis, *Arthrobacter* sp. α11c was selected for genome sequencing, as it showed the highest growth potential on α-Hed and HedC, together with *Pseudoarthrobacter* α12b, a closely related strain not able to grow on any of the compounds tested.

Genome analysis revealed that *Arthrobacter* sp. α11c carries 47 putative glycosidase genes belonging to 25 subclasses. In comparison, *Pseudoarthrobacter* α12b carries 40 putative glycosidase genes belonging to 19 subclasses (**Fig. 2**). Comparative genome analysis between *Arthrobacter* α11c and *Pseudoarthrobacter* α12b revealed that *Arthrobacter* sp. α11c harbors 14 glycosidases that are not present in *Pseudoarthrobacter* sp. α12b (**Fig. 2**). Specifically, we identified a β-glucosidase (*bglB1*; LOFIDKPF_00997; EC. 3.2.1.21) and two β-galactosidases (LOFIDKPF_01064 and LOFIDKPF_01065; EC. 3.2.1.23) which are not present in the genome of *Pseudoarthrobacter* α12b (**Fig. 2**).

**Figure 2.**
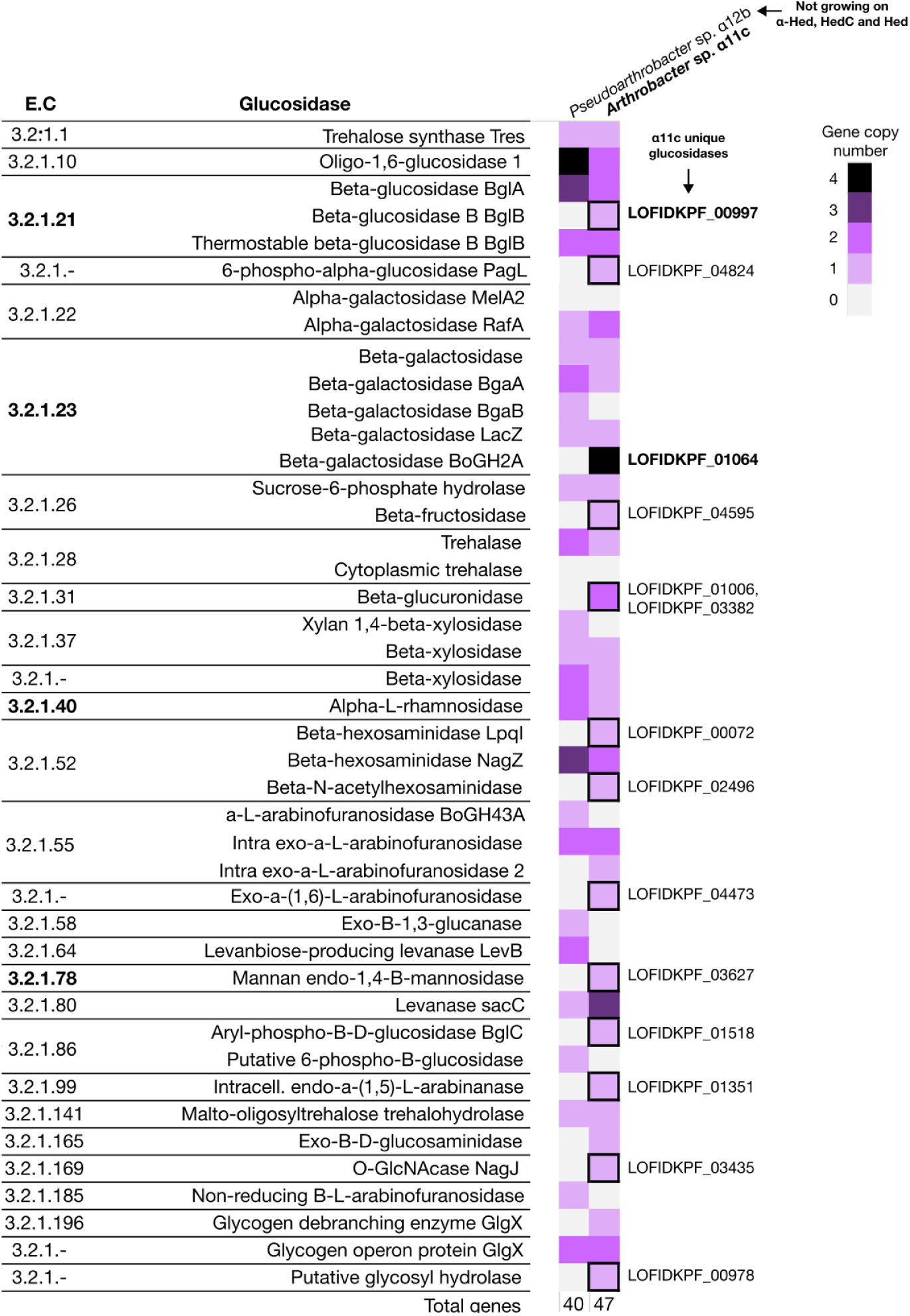
Glycosidases putatively involved in saponin turnover. Glucosidases (EC 3.2.1) from the genome of *Arthrobacter* sp. α11c, which can grow on saponins, as well as from *Pseudoarthrobacter* sp. α12b, which cannot grow on saponins were identified based on Prokka (Version 1.13) annotations (Seemann, 2014), and filtered using the SeqIO module of BioPython (Cock et al., 2009). β-glucosidases (EC 3.2.1.21), β-galactosidases (EC 3.2.1.23), and α-L-rhamnosidase (EC 3.2.1.40), which have been reported to deglycosylate saponins, are shown in bold. The heat-map represents the copy number of genes from each glucosidase identifier. Unique glucosidases from *Arthrobacter* sp. α11c are labeled (LOFIDKPF_).

Comparative genome analysis of *Arthrobacter* sp. α11c and *Pseudoarthrobacter* α12b was done to search for glycosidase genes in the unlinked regions, *i.e.,* regions exhibiting <5% sequence identity between the genomes over a minimum of 1000 bp. The position of the largest unlinked region spanning ≈250 Kbp, is indicated in **Fig. 3a**. In this unlinked region, a glycosidase gene cluster including an α-L-mannosidase (EC 3.2.1.40) and the β-glucosidase (*bglB1*; LOFIDKPF_00997; EC 3.2.1.21) was found on the *Arthrobacter* α11c genome. Within the same cluster, a gene encoding a 3α,20β-hydroxysteroid dehydrogenase (3α,20β-HSD; EC 1.1.1.53) was identified (**Fig. 3b**). Additionally, we found a second glycosidase gene cluster in the same unlinked region containing two β-galactosidase-encoding genes (LOFIDKPF_01064 and LOFIDKPF_01065; 3.2.1.23), as well as a 3-oxo-Delta(4)-steroid 1-dehydrogenase encoding gene (KSTD; EC 1.3.99.4) (**Fig. 3c**)(**Table S1**). No glycosidase genes were found on the corresponding unlinked region (≈930-1150kb) in the *Pseudoarthrobacter* α12b genome (**Fig. S3**).

**Figure 3.**
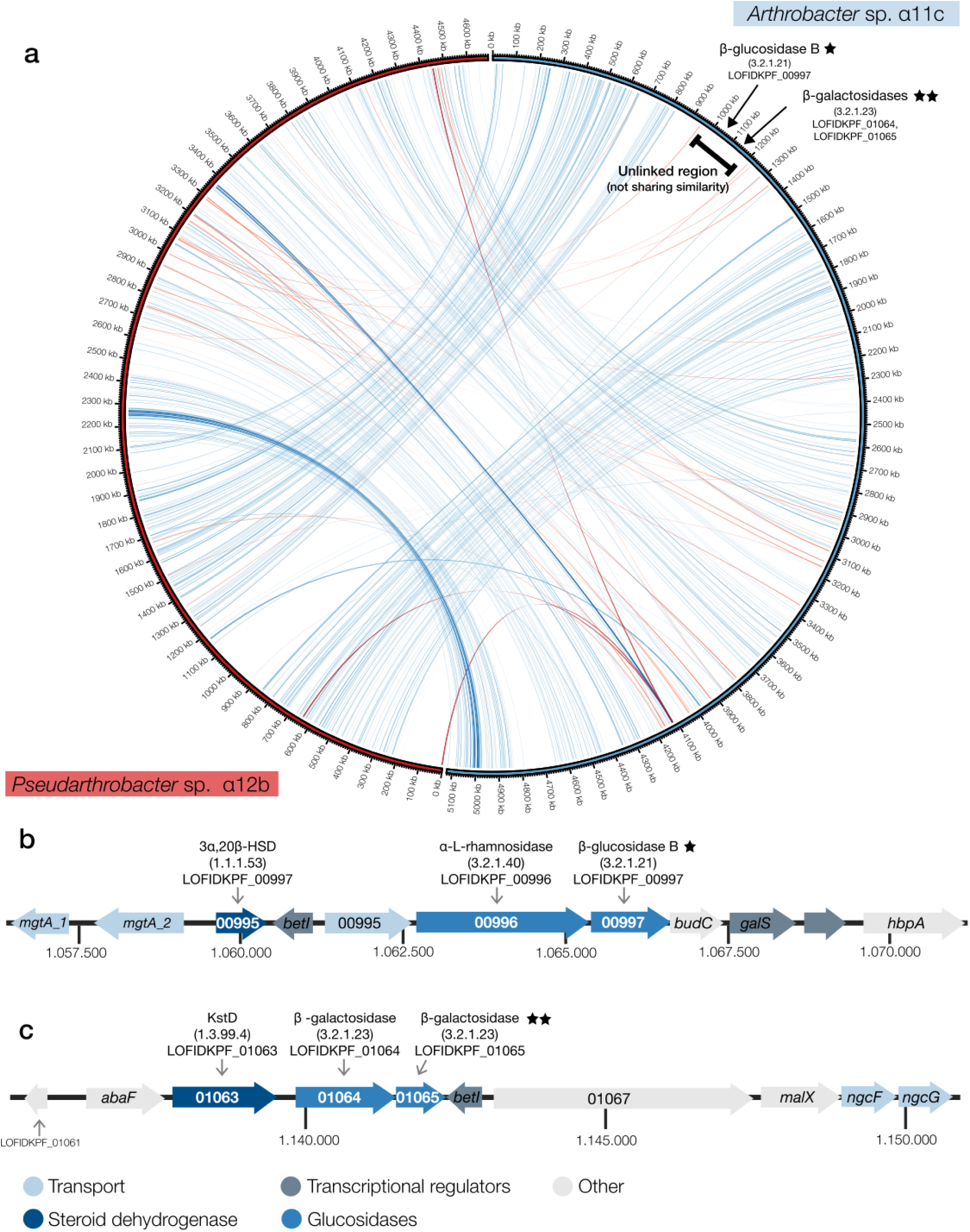
Glycosidase containing clusters identified in *Arthrobacter* sp. α11c. **a)** Complete genome alignment between *Arthrobacter* sp. α11c that can grow on saponins and the closely related *Pseudoarthrobacter* sp. α12b that cannot grow on saponins. Genome alignment and visualization were performed by MUMer2Circos (Version 1.4.2) (https://github.com/metagenlab/mummer2circos). Linked regions have a sequence identity higher than 95% and a length higher than 1000bp. Blue lines represent aligned segments that are in the same orientation with respect to the query (*Arthrobacter* sp. α11c), and red lines represent aligned segments in reversed orientation to the query (*Arthrobacter* sp. α11c). Glucosidases from *Arthrobacter* sp. α11c belonging to the main putative saponin deglycosylating enzymes (EC 3.2.1.21 and EC 3.2.1.23) are shown in the genome map with an arrow. **b)** and **c)** Genomic context of the *Arthrobacter* sp. α11c clusters from the unlinked region harboring glucosidases (α-L-rhamnosidase: EC 3.2.1.40, β-glucosidase EC 3.2.1.21, and β-galactosidase EC 3.2.1.23) and steroid-degrading enzymes (3α,20β-HSD: EC 1.1.1.53 and KstD: EC 1.3.99.4). In **b)** Adjacent genes are *mgtA_2*: Magnesium-transporting ATPase; LOFIDKPF_00993: 3α-(or 20β)-hydroxysteroid dehydrogenase; *betI_6*: HTH-type transcriptional regulator BetI (TetR/AcrR family); LOFIDKPF_00995: Putative diacylglyceride transporter/D-galactonate/Major Facilitator Superfamily (MFS); LOFIDKPF_00996: α-L-rhamnosidase; *bglB_1*: β-glucosidase B; *budC*: Diacetyl reductase (S)-acetoin forming; *galcS*: HTH-type transcriptional regulator GalS; LOFIDKPF_00100: Hypothetical protein (TetR-family transcriptional regulator); In **c)** LOFIDKPF_01061: Hypothetical protein; *abaF*: Fosfomycin resistance protein; *kstD*: 3-oxosteroid 1-dehydrogenase; LOFIDKPF_01064: β-galactosidase BoGH2A; LOFIDKPF_0165: β-galactosidase BoGH2A; *betI_8*: HTH-type transcriptional regulator BetI (TetR/AcrR family); LOFIDKPF_01067: Hypothetical protein; *malX_1*: Maltose/maltodextrin-binding protein; *ngcF_2*: Diacetylchitobiose uptake system permease protein.

### Glucosidase activity

To demonstrate the presence of glucosidase activity in *Arthrobacter* sp. α11c, we used the pNPG colorimetric method on cultures grown with α-Hed and HedC. The results showed the presence of glucosidase activity on both saponins; however, this activity was not significantly higher than control samples (**Fig. 4**). The results indicate that the glucosidase activity in *Arthrobacter* sp. α11c is not specifically induced by saponins.

**Figure 4.**
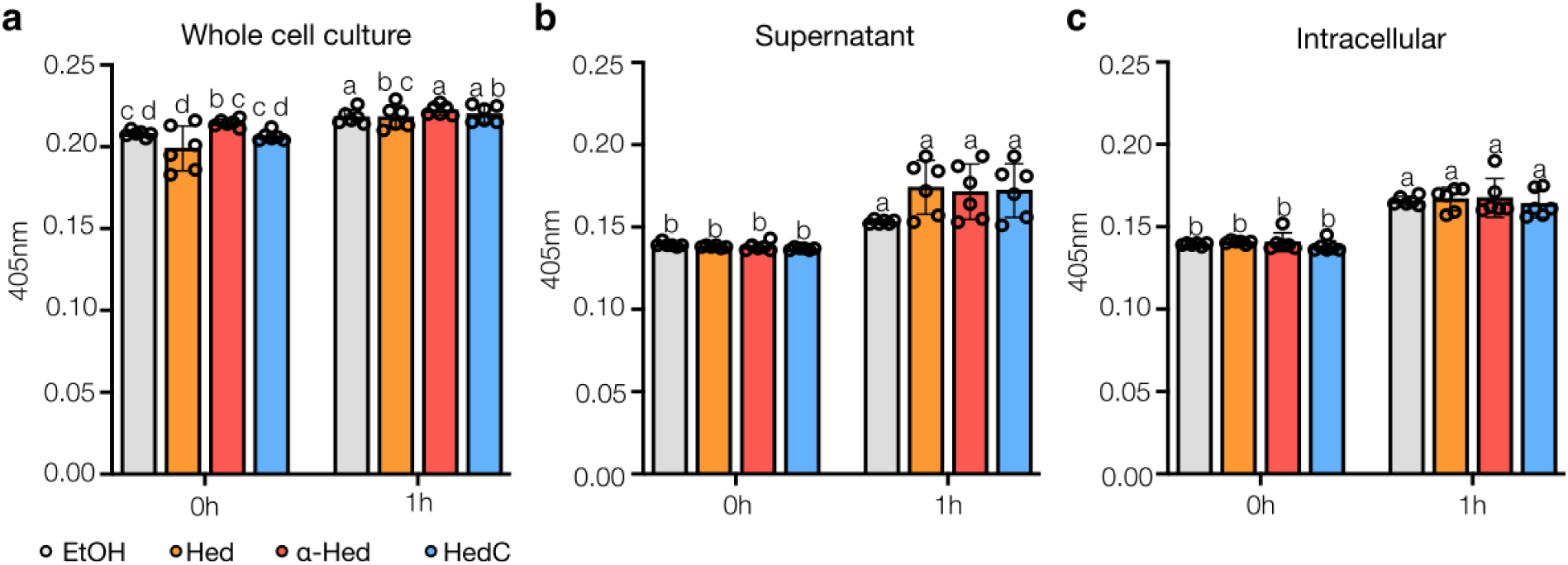
Glucosidase activity from *Arthrobacter* sp. α-11c. Glucosidase activity from **a)** whole cell culture, **b)** culture supernatant, and **c)** intracellular fraction of *Arthrobacter* sp. α11c. *Arthrobacter* sp. α11c cells were inoculated into media containing saponins (α-Hed and HedC) or the corresponding sapogenin (Hed) to a final concentration of 20 µM, each dissolved in 1% EtOH (used as control), and harvested after 24h of incubation. Glucosidase activity was performed using the colorimetric method with pNPG (5 mM). Open circles represent individual values from biological replicates. Statistical test, ordinary two-way ANOVA.

### Saponin degradation by Arthrobacter sp. α11c

To verify degradation of saponins by *Arthrobacter* sp. α11c, we incubated cultures with α-Hed, HedC, and Hed and monitored their concentrations extracellularly and intracellularly after 0, 2, and 24 hours as a proxy for degradation. To specifically distinguish between extracellular degradation, and intracellular degradation, the cultures were separated into supernatant, ethanol cell-washed, and intracellular fractions. The results showed a significant decrease in the concentrations of α-Hed, HedC, and Hed in the supernatant, with reductions of 95%, 91%, and 98%, respectively (**Fig. 5ace**). No residual saponins were found in the ethanol-washed fraction, which rules out adsorption to the cells as the primary cause of concentration reduction in the supernatant. Additionally, the intracellular measurements confirmed the uptake of α-Hed, HedC, and Hed, with a peak observed in the 2h samples. Taken together, the results indicate that saponins and Hed are taken up and intracellularly metabolized by *Arthrobacter* sp. α11c. No apparent degradation compounds were detected for any of the two tested saponins, as confirmed by both gas chromatography (GC) and liquid chromatography (LC) (**Fig. 5bdf**). To evaluate whether the saponins were degraded into their sapogenin (Hed) backbone, we used GC-MS to specifically search for the characteristic 203 m/z fragment (Santos et al., 2018), which is predicted to arise from silylated derivatives of α-Hed and HedC after enzymatic cleavage to Hed. However, no peaks corresponding to this fragment were detected under the applied conditions. These findings suggest that not only the saponins, but also the sapogenin itself, were completely degraded.

### Degradation kinetics

To investigate the intracellular uptake and degradation of saponins by *Arthrobacter* sp. α11c, saponin and sapogenin levels in the supernatant and intracellular fraction were measured over a six-hour period. The cell-washed fraction was excluded from this assay, as no precipitated saponins were detected in this fraction (**Fig. 5**). The highest α-Hed intracellular concentration was observed at 40 minutes after inoculation (**Fig. 6a**), and the highest intracellular concentration of HedC was observed at 60 minutes after inoculation (**Fig. 6b**). In contrast, Hed was found intracellularly immediately after inoculation. In agreement with the intracellular transport, the half-life of HedC was the highest (86 minutes), followed by α-Hed (59 minutes) and Hed (15 minutes) (**Table 1**). Full degradation of α-Hed and HedC was observed after 6 h of incubation (**Fig. 6ab**), whereas the sapogenin Hed was fully degraded within one hour of incubation (**Fig. 6c**).

**Figure 5.**
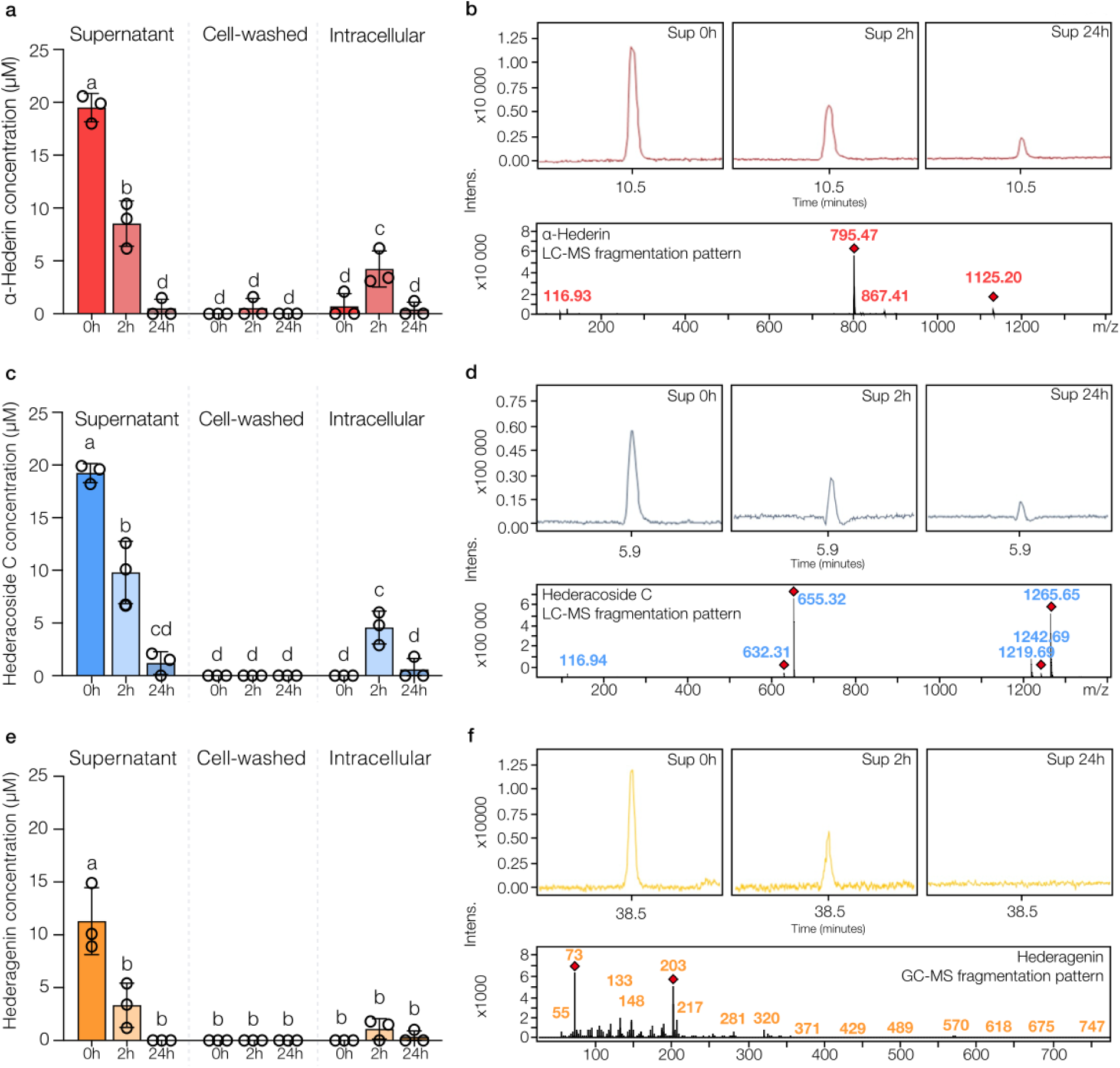
Uptake and degradation of saponins by *Arthrobacter* sp. α**11c.** Temporal concentration profiles of the two saponins and the sapogenin hederagenin (Hed) **a)** and **b)** LC-MS analysis; α-Hed (red), LC-MS analysis; 795.47 m/z corresponds to the formic acid adduct of α-hederin. **c)** and **d)** Hed C (blue), 1265,65 m/z corresponds to the formic acid adduct of Hed C, the 655,32 m/z to the Hed C that has lost 4 sugars. **e)** and **f)** GCMS analysis of sapogenin Hed (orange) in *Arthrobacter* sp. α11c cultures. Concentrations were measured at 0, 2, and 24 hours in the supernatant, cell-washed fraction, and intracellular fraction using LC-MS for α-Hed and HedC, and GC-MS for sapogenin Hed. Mass spectrometric analysis confirmed compound identity and monitored potential degradation products. The 203 m/z corresponds to characteristic degradation product of hederagenin under GCMS analysis (Santos et al., 2018) All data represent mean ± SD from three biological replicates. Different letters indicate statistically significant differences (two-way ANOVA, p < 0.05). α-Hed and HedC were detected by monitoring their sodium adducts, with m/z values of 795.5 ± 0.05 and 1265.5 ± 0.05, respectively. HedC, being more hydrophilic, eluted at 5.9 minutes, while α-Hed eluted at 10.5 minutes.

**Figure 6.**
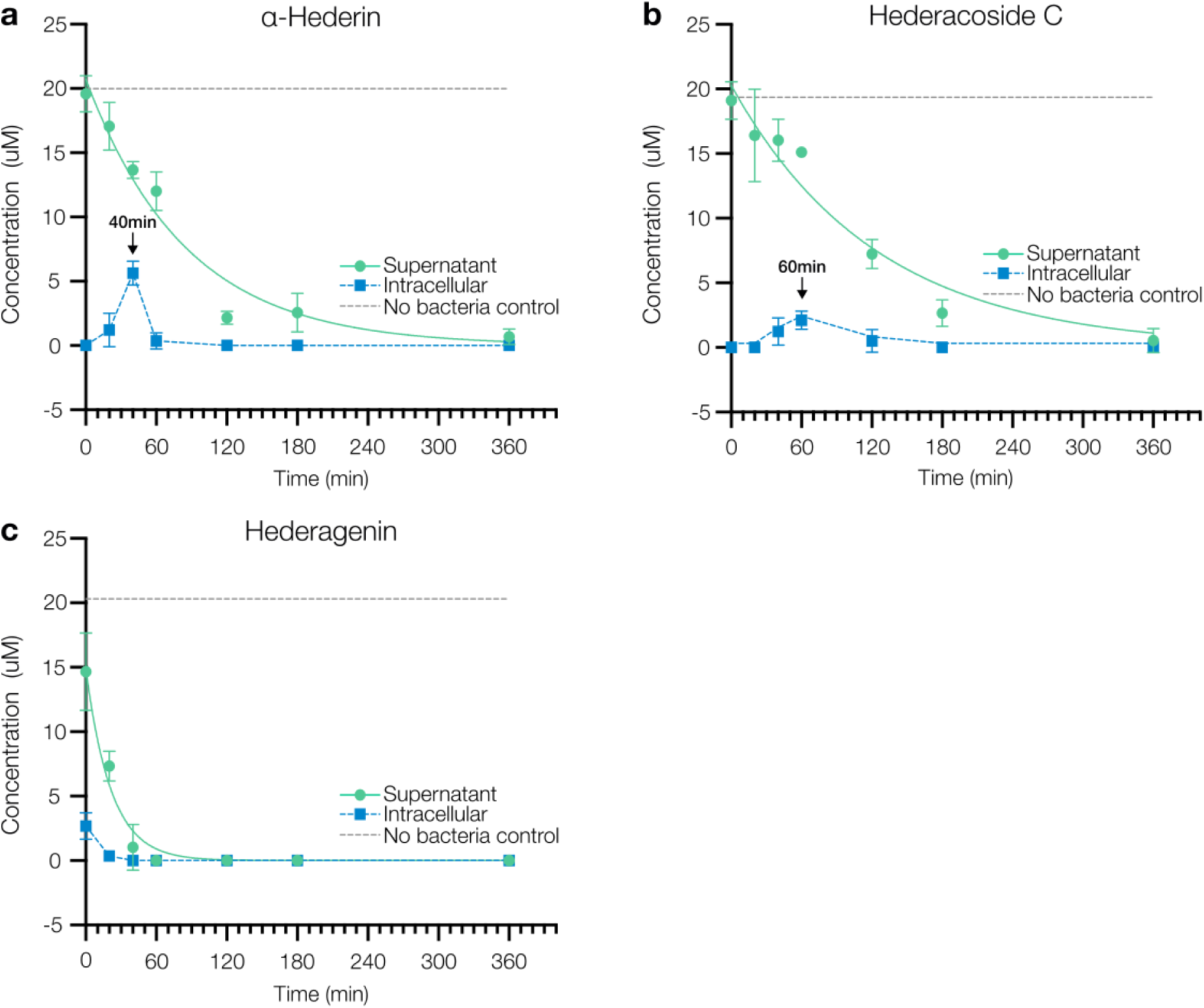
Saponin intracellular uptake and degradation rate by *Arthrobacter* sp. α-11c. Intracellular uptake and degradation rates of **a)** α-Hed and **b)** HedC saponins, and **c)** Hed sapogenin. The decrease in α-Hed, HedC, and Hed concentrations in the supernatants follows first-order decay kinetics and is represented by solid green lines, while dashed blue lines indicate intracellular concentrations for visual guidance. The initial concentration of α-Hed, HedC, and Hed was 20 µM in 1% EtOH, and measured concentrations are shown as dashed gray lines. *Arthrobacter* sp. α11c was inoculated at OD₆₀₀ = 0.1. Samples were collected after 20, 40, 60, 120, 180, and 360 minutes of incubation. Saponin concentrations were quantified by LC-MS, and sapogenin concentrations by GC-MS. Saponin and sapogenin concentrations are presented as mean ± standard deviation from three biological replicates.

**Table 1.** Degradation kinetics of hederagenin-based saponins by *Arthrobacter* sp. α11c.

| | α-Hed | HedC | Hed |
| --- | --- | --- | --- |
| Initial concentration (μM) | 20.7 | 20.3 | 15.0 |
| Decay Constant $k$ ($\text{min}^{-1}$) | $0.01177 \pm 0.0015$ | $0.008062 \pm 0.0006$ | $0.04663 \pm 0.0122$ |
| Half-life (min) | 59 | 86 | 15 |
| $R^2$ | 0.9421 | 0.9138 | 0.9342 |

## Discussion

Microorganisms possess enzymatic systems capable of breaking down complex chemical structures, which play a crucial role in mitigating the environmental risks associated with e.g. pesticides. Saponins have been proposed as potential biopesticides; however, it is essential to assess whether soil microorganisms can biodegrade saponins to prevent their persistence in the environment and ensure their safe application on crops. While previous studies have demonstrated that soil bacteria can degrade steroidal saponins, there is currently no published evidence of full biodegradation of triterpenoid saponins and sapogenins.

In this study, we show that two hederagenin-derived triterpenoid saponins, α-hederin (α-Hed) and hederacoside C (HedC) can be degraded by *Arthrobacter* sp. Α11c as confirmed by LCMS and GCMS analysis (**Fig. 5 and Fig. 6**). The transient presence of saponins in the intracellular fraction, together with the decrease in concentration in the supernatant, and no detection of degradation products by LCMS or GCMS, suggests that α-Hed and HedC are transported into the bacterial cells for degradation to support growth. Although the sapogenin hederagenin (Hed) was rapidly internalized and degraded, no degradation products were detected via LCMS or GCMS analysis. Moreover, Hed did not promote significant growth of *Arthrobacter* sp. α11c. One explanation could be the high metabolic cost associated with its degradation due to the lack of sugar, or the formation of non-utilizable byproducts. Alternatively, Hed may have been sequestered in the final pellet during sample processing. However, this scenario appears unlikely, given its absence of detection in the ethanol wash and the intracellular fraction.

To obtain insight into the potential glycosidases involved in saponin degradation, we performed a comparative genome analysis of *Arthrobacter* sp. α11c and the closely related *Pseudoarthrobacter* sp. α12b, which does not grow on α-Hed, HedC nor the sapogenin Hed. This revealed that *Arthrobacter* sp. α11c possesses a high diversity of glycosidases. Furthermore, the annotation showed 14 glycosidases identified in *Arthrobacter* sp. α11c which are not found in *Pseudoarthrobacter* sp. α12b. Examination of the identified unlinked region containing putative glycosidases identified glycosidase homologs (EC 3.2.1.21, 3.2.1.23, and 3.2.1.40) previously reported for saponin deglycosylation (Hennessy et al., 2020;Park et al., 2017; Wang et al., 2022). Moreover, these unique clusters present in *Arthrobacter* α11c also contain the 3α,20β-HSD and KstD steroid-dehydrogenase coding genes previously found in steroid-degrading bacteria (Galán et al., 2017). The localization of 3α,20β-HSD and KstD coding genes within the glycosidase gene clusters suggests a potential role for these steroid-degrading enzymes in the observed saponin degradation.

The only reported sapogenin biotransformation is, to the best of our knowledge, from the Actinobacterium *R. rhodochrous* IEGM 66, which converted betulin into betulone (Grishko et al., 2013; Maltseva et al., 2024). The biotransformation of betulin to betulone is achieved by oxidizing the 3α-hydroxyl group to the corresponding ketone. Hence, the 3α,20β-HSD may encode for enzymes that perform such a transformation in both the betulin and the sapogenin Hed, with KstDs performing the subsequent step of A-ring degradation (Wójcik et al., 2023). Bacteria containing enzyme homologs for these two steps have also been reported in bacteria performing steroid degradation (Petrusma et al., 2014; Wójcik et al., 2023). The KstD found in *Arthrobacter* sp. α11c showed homology (38% a.a. seq. id) with the KstD (7P18) from *Sterolibacterium denitrificans* Chol-1, which is a promiscuous (Wójcik et al., 2023). These findings suggest that KstD from *Arthrobacter* sp. α11c may be involved in the A-ring oxidation of hederagenin-based saponins.

Glucosidase activity was confirmed in *Arthrobacter* sp. α11c in the supernatant and in the intracellular culture fractions. Cells cultured with the saponins α-Hed and HedC as well as the sapogenin Hed showed no significant difference in glucosidase activity compared to cells grown in the absence of saponins (**Fig 4**). This indicates that the saponins used in this study do not induce the basal glucosidase activity of *Arthrobacter* sp. α11c. Future studies are needed to reveal the regulatory network behind saponin degradation in *Arthrobacter* sp. α11c.

In conclusion, our findings demonstrate that hederagenin-based saponins can be fully degraded by *Arthrobacter* sp. Α11c, and that this bacterium can use saponins as a carbon source to support growth. The results indicate that saponins are taken up intracellularly, followed by deglycosylation and complete degradation of the sapogenin, demonstrating the ability of soil microorganisms to fully and quickly degrade the hederagenin-based saponins. This knowledge is important for the risk-assessment of saponin-based bioinsecticides.

## Declarations

### Ethics approval and consent to participate

Not applicable

### Consent for publication

Not applicable

## Availability of data and materials

The 16S rRNA genes of the strain library isolates are available in Genbank under accession numbers PV902503-PV902532. The fully assembled genomes are available under BioProject PRJNA1288778. All other raw data are available on request.

## Authors’ contributions

CW: Investigation, Writing - review & editing.

HCBH: Conceptualization, Supervision, Funding acquisition, Project Administration, Resources, Writing - review & editing

KYMZ: Conceptualization, Formal analysis, Investigation, Software, Visualization, Writing – original draft, Writing - review & editing.

MD: Conceptualization, Formal analysis, Investigation, Visualization, Writing – original draft, Writing - review & editing.

MHN: Conceptualization, Supervision, Funding acquisition, Project Administration, Resources, Writing - review & editing

MR: Investigation. PDB: Investigation.

SB: Conceptualization, Supervision, Funding acquisition, Project Administration, Resources, Writing - review & editing

## Acknowledgments

Not applicable

