## Supplementary figures and Tables for "Degradation of biopesticidal triterpenoid saponins by the soil bacterium *Arthrobacter* sp. α-11c"

### Supplementary Table and Figures

**Table S1.** Candidate genes for steroid degradation pathway enzymes identified in *Arthrobacter* sp. α11c

| Short name | Protein | E.C. | Reported homologs | Locus tag | a.a. sequence similarity % | Reference |
|---|---|---|---|---|---|---|
| 3α,20β-HSD | 3α-(or 20β)-hydroxysteroid dehydrogenase | 1.1.1.53 | 1HDC | LOFIDKPF_00993 | 50.42 | (Ghosh et al., 1994; Ghoshl et al., 1994) |
| KstD | 3-oxosteroid 1-dehydrogenase | 1.3.99.4 | 7P18 | LOFIDKPF_01063<br>LOFIDKPF_01337 | 38.43<br>39.58 | (Wójcik et al., 2023). |
| KshA | 3-ketosteroid-9-alpha-monooxygenase, oxygenase component | 1.14.15.30 | 4QCK | --- | --- | (Capyk et al., 2011) |
| KshB | 3-ketosteroid-9-alpha-monooxygenase, ferredoxin reductase component | 1.14.15.30 | P9WJ93 | --- | --- | (Capyk et al., 2009) |
| HsaA | Flavin-dependent monooxygenase, oxygenase subunit HsaA | 1.14.14.12 | Q0S811<br>5lvu | LOFIDKPF_01105 | 36.46 | (Kugel et al., 2017; Van der Geize et al., 2007) |
| HsaC | Iron-dependent extradiol dioxygenase | 1.13.11.25 | Q9KWQ5 | --- | --- | (Van der Geize et al., 2007) |
| HsaD | 4,5:9,10-diseco-3-hydroxy-5,9,17-trioxoandrosta-1(10),2-diene-4-oate hydrolase | 3.7.1.17 | Q9KWQ6 | --- | --- | (Van der Geize et al., 2007) |

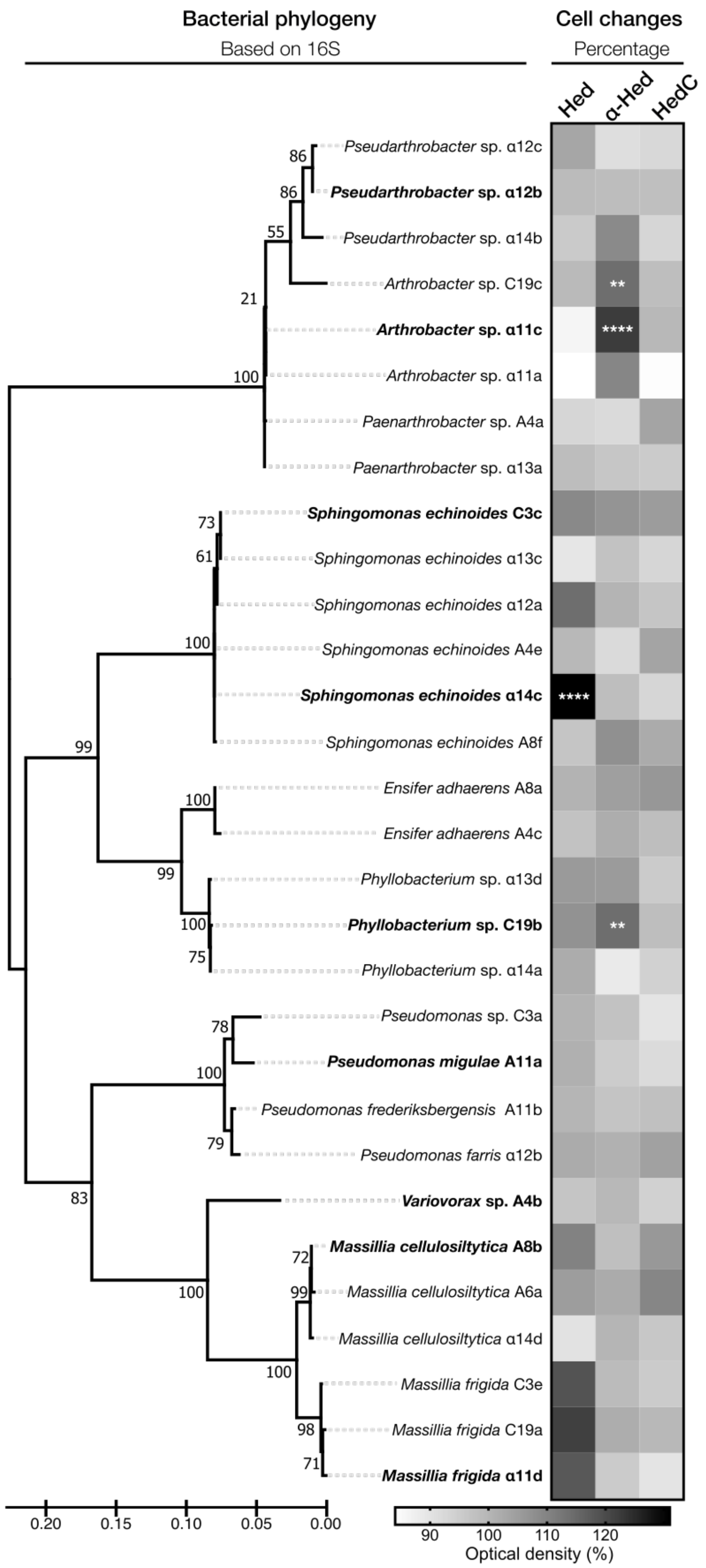

Bacterial phylogeny
Based on 16S
Cell changes
Percentage
Hed
α-Hed
HedC
Pseudarthrobacter sp. α12c
Pseudarthrobacter sp. α12b
Pseudarthrobacter sp. α14b
Arthrobacter sp. C19c
Arthrobacter sp. α11c
Arthrobacter sp. α11a
Paenarthrobacter sp. A4a
Paenarthrobacter sp. α13a
Sphingomonas echinoides C3c
Sphingomonas echinoides α13c
Sphingomonas echinoides α12a
Sphingomonas echinoides A4e
Sphingomonas echinoides α14c
Sphingomonas echinoides A8f
Ensifer adhaerens A8a
Ensifer adhaerens A4c
Phyllobacterium sp. α13d
Phyllobacterium sp. C19b
Phyllobacterium sp. α14a
Pseudomonas sp. C3a
Pseudomonas migulae A11a
Pseudomonas frederiksbergensis A11b
Pseudomonas farris α12b
Variovorax sp. A4b
Massillia cellulosiltytica A8b
Massillia cellulosiltytica A6a
Massillia cellulosiltytica α14d
Massillia frigida C3e
Massillia frigida C19a
Massillia frigida α11d
0.20 0.15 0.10 0.05 0.00
90 100 110 120
Optical density (%)

**Figure S1. Bacterial growth of 30 soil bacterial isolates cultured in media supplemented with the saponins sapogenin.** In bold are highlighted the nine strains that based on increase in OD600 and taxonomic diversity were selected for further analysis. The maximum likelihood phylogenetic tree was constructed based on a multiple alignment of 16S rRNA gene sequences. Bootstraps values (1000 iterations) are shown as at the branch nodes. Bacterial cells were inoculated in media supplemented with saponins (α-Hed or HedC) and their corresponding sapogenin (Hed) at a final concentration of 20 μM dissolved in EtOH 1%. The final OD was measured after 24 hours of incubation. Data was collected from three biological replicates. Statistical test, two-way ANOVA; ns, not significant; **, $P$<0.01, ****, $P$< 0.0001.

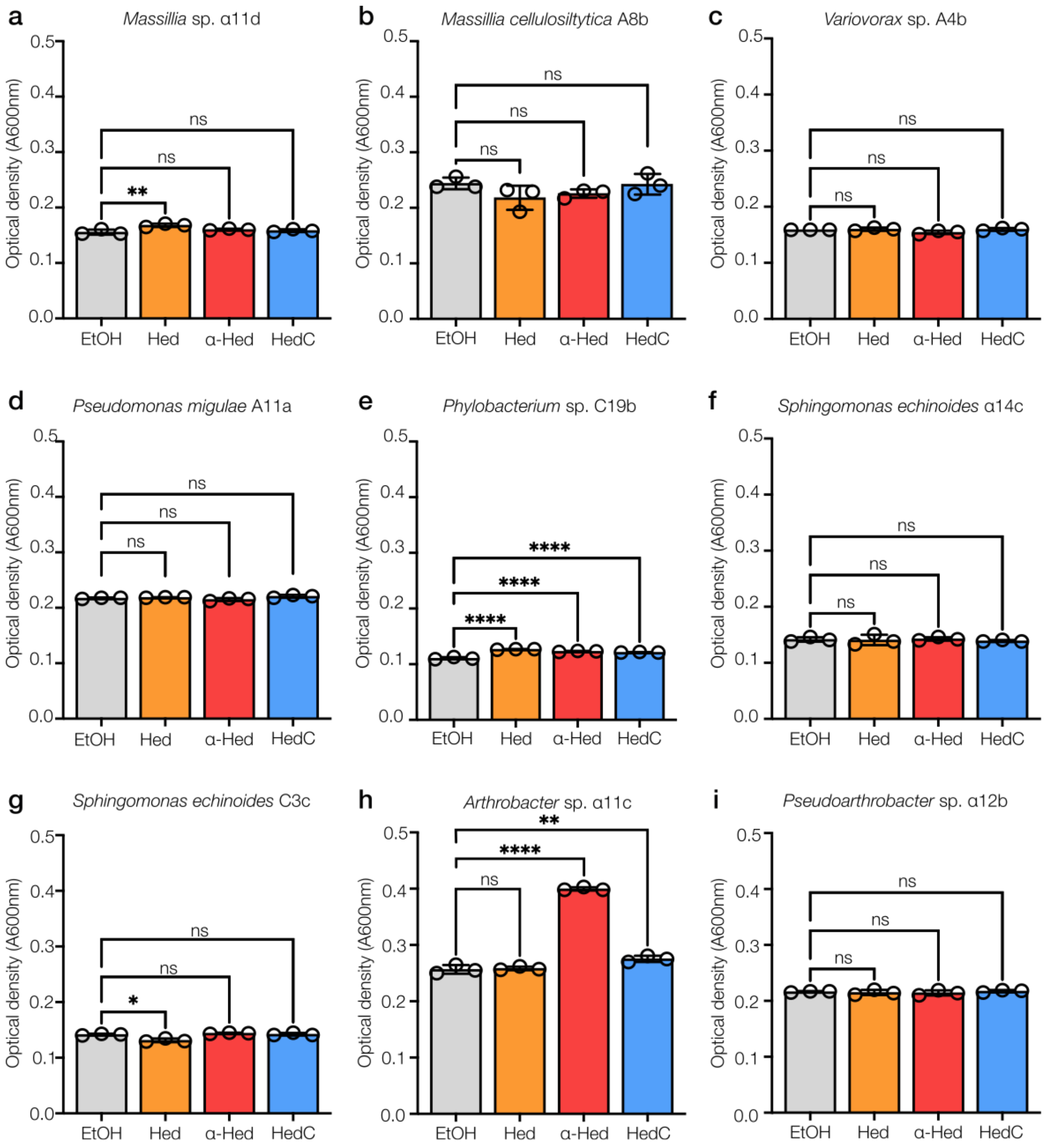


**Figure S2. Increase in OD600 for the nine selected soil bacterial isolates in media supplemented with saponins or sapogenin.** Bacterial cells were inoculated with the glycosylated saponins (α-Hed and HedC) and their sapogenin (Hed) at a final concentration of 20 μM in EtOH 1%, controls contained only EtOH 1%. Final OD600 was measured after 24h of incubation. Statistical test, one-way ANOVA; ns, not significant; **, $P$<0.01, ****, $P$< 0.0001. Data was collected from three biological replicates

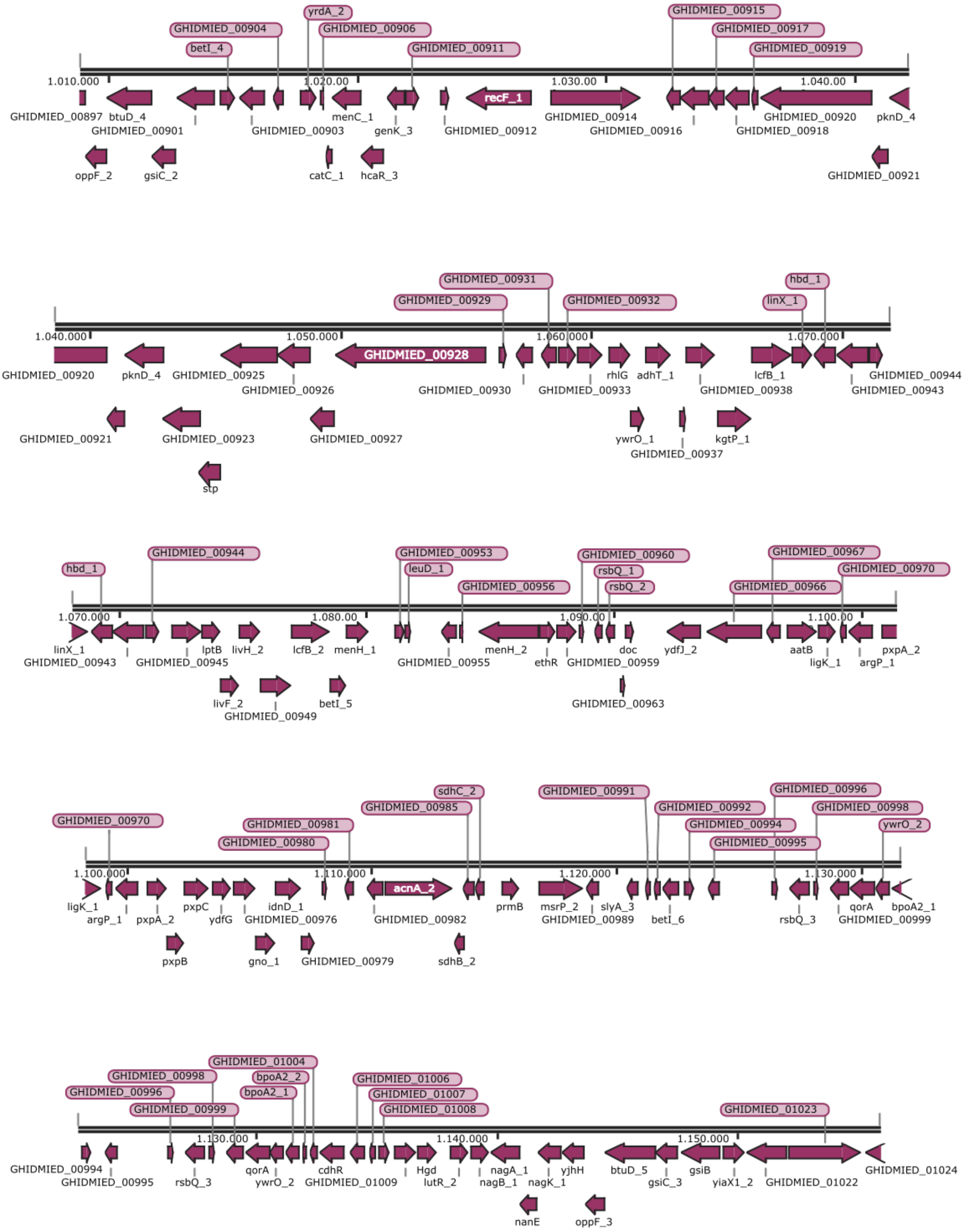


**Figure S3. Annotation of the unlinked region in *Pseudoarthrobacter* sp. α12b genome.** No glucosidases nor putative steroid degrading enzymes from the glucosidase clusters identified in the unlinked region of the *Arthrobacter* α11c genome were found in the unlinked region of the *Pseudoarthrobacter* α12b genome.

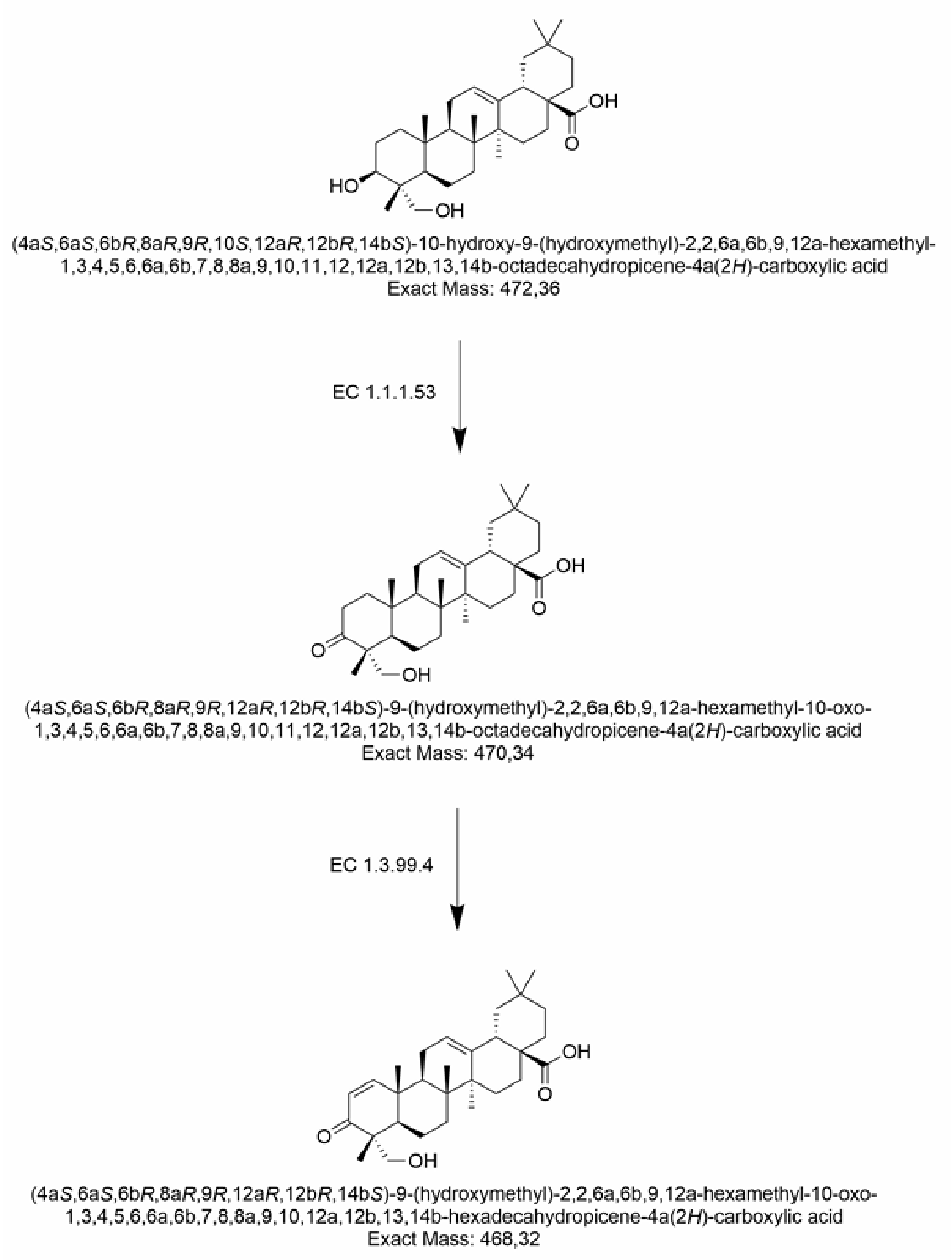


**Figure S4. Putative degradation steps catalyzed by steroid-degrading enzymes found in *Arthrobacter* sp. α11c.** The first step, the A-ring oxidation, is predicted to be performed by 3α,20β-HSD (EC 1.1.1.53), followed by KstD (EC 1.3.99.4), resulting in a modified hederagenin of 468.32 WM.

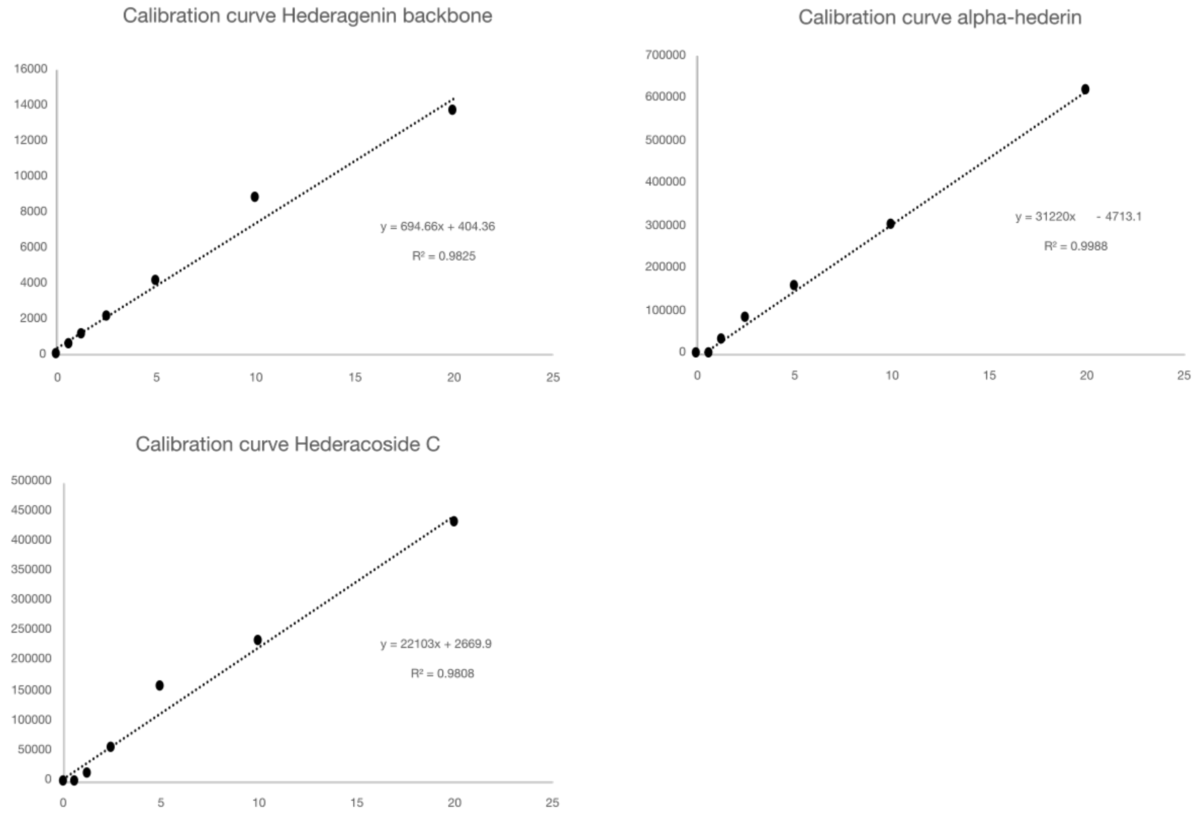


**Figure S5. Calibration curves used to calculate the concentration of saponins (α-Hed and HedC; LCMS) and sapogenin (Hed; GCMS).**
